# Characterization of the HHV-6 U20 immunoevasin

**DOI:** 10.1101/2022.10.10.511687

**Authors:** Christine L. Schneider, Melissa L. Whyte, Sheryl L. Konrad, Amy W. Hudson

**Affiliations:** Department of Microbiology and Immunology, Medical College of Wisconsin, Milwaukee, WI, USA

**Keywords:** U20, HHV-6A, HHV-6B, HHV-7, herpesviruses, immune evasion

## Abstract

Roseoloviruses (HHV-6A, -6B, and -7) infect >90% of the human population during early childhood, and are thought to remain latent or persistent throughout the life of the host. As such, *these viruses are among the most pervasive and stealthy of all viruses;* they must necessarily excel at escaping immune detection throughout the life of the host, and yet very little is known about how these viruses so successfully escape host defenses. Herein, we characterize the HHV6A and HHV6B U20 gene products, which are encoded within a block of genes unique to the roseoloviruses, and therefore of particular interest. Despite 92% amino acid identity, U20 proteins from HHV6A and 6B have been shown to possess different host evasion functions. Here we characterize expression, trafficking, and post-translational modifications of U20 during HHV6A infection. While U20 localized to lysosomes in HHV-6A-infected cells, HHV-6B U20 trafficked to the cell surface and was rapidly internalized. HHV-6B U20 trafficked slowly through the secretory system, receiving several post translational modifications to its N-linked glycans indicative of surface expressed glycoproteins. Interestingly, U20 is also phosphorylated on at least one Ser, Thr, or Tyr residue. These results provide a framework to understand the role(s) of U20 in evading host defenses.

**Importance:** HHV6A and HHV6B U20 proteins are virus-encoded integral membrane glycoproteins possessing class I MHC-like folds. As such, it is tempting to speculate that they are involved in host evasion. Indeed, although they share 92% identity, HHV6A U20 has been shown to target NK activating ligands (1) and HHV6B U20 has been shown to inhibit TNF receptor signaling and apoptosis (2). Here, we have performed cell biological and biochemical characterization of the trafficking, glycosylation, and post-translational modifications occurring on HHV6A and -6B U20, and we demonstrate U20 expression in the context of HHV6 infection.

## Introduction

Herpesviruses employ a wide variety of strategies to thwart host anti-viral responses. In establishment and maintenance of lifelong infections in immunocompetent hosts, the herpesviruses must continuously navigate interactions with host anti-viral countermeasures. As part of their arsenal, herpesviruses encode proteins that interfere with class I MHC presentation, presumably to protect the virus infected cell from recognition by cytotoxic T cells (for review see (3)). Herpesviruses also encode proteins that interfere with NK cell recognition (for review see (4)). Importantly, characterization of proteins from these viruses have proven valuable in expanding our understanding of fundamental cellular processes like antigen presentation, ER-associated degradation, and NK cell recognition (5–13).

Human cytomegalovirus (HCMV) is the standard-bearer in this arena of herpesvirus host evasion; 4 HCMV ORFs have been shown to downregulate class I MHC molecules, and 5 HCMV ORFs and an HCMV-encoded microRNA have been shown to downregulate NK activating ligands (Reviewed in (14)). In contrast to HCMV, there are relatively few studies characterizing proteins from the roseoloviruses (HHV-6A, -6B, and -7). HHV-6 and 7 lack positional homologs of HCMV host evasion proteins, and instead contain several blocks of genes unique to the roseoloviruses, some of which have been shown to encode proteins involved in evasion of host defenses (15–20). For example, the U21 open reading frame from HHV-6A, 6B and 7 is a type I integral membrane protein that reroutes class I MHC to lysosomes or degradation (15, 18, 21). The same U21 protein has been shown to downregulate NK activating ligands and prevent NK killing of U21-expressing cells (16). U24 from HHV-6A, 6B and 7 is a tail-anchored protein whose expression results in reduced surface expression of T cell receptors (19, 20). U26 from HHV-6B inhibits innate antiviral responses by interfering with MAVS signaling (22). Surprisingly, although U20 from HHV-6A and -6B share 92% identity (Figure 1), recent studies ascribe different functions to HHV6A U20 and HHV6B U20. HHV6A U20 was shown to downregulate NKG2D ligands, while HHV6B U20 was shown to inhibit TNFα-induced apoptosis during nonproductive infection with HHV6B (1, 2).

**Figure 1.**
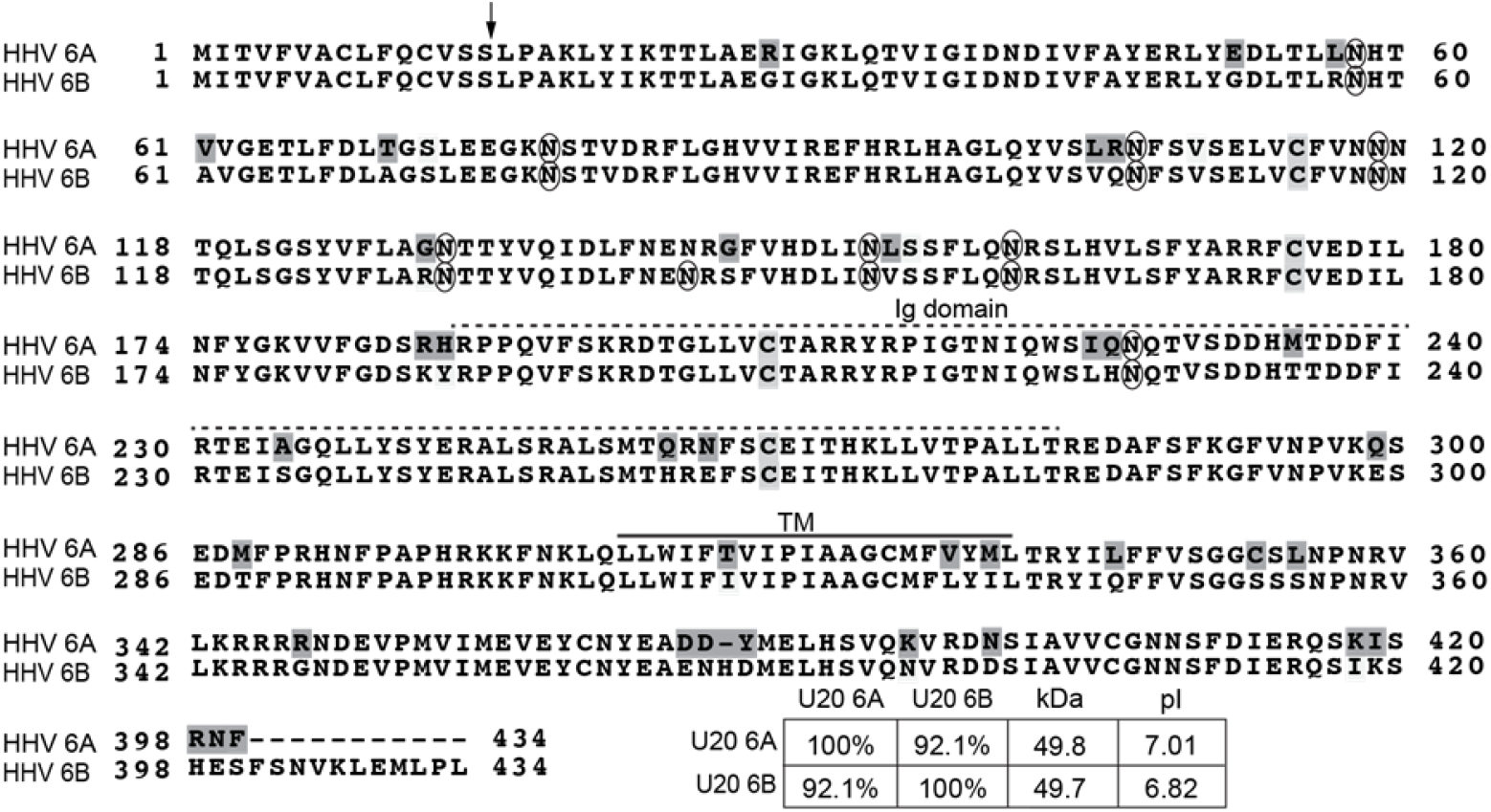
Alignment HHV6A and -6B U20. Amino acid residues that differ between HHV6A and -6B are highlighted in dark gray. Conserved cysteine residues are highlighted in light gray. The arrow depicts the predicted signal peptide cleavage site. Predicted immunoglobulin domains, transmembrane domains (solid line), and glycosylation sites (circles) are noted. The inset table indicates the percent identity between U20 proteins from HHV6A and 6B as well as predicted sizes and isoelectric points.

Several features of U20 make it attractive as a potential host-defense evasin: like other host-defense evasive proteins encoded by HCMV, *U20* is located within a block of other host-evasive genes, and it is expressed with early kinetics(23, 24). Results from recent structural modeling studies suggest that U20 contains an immunoglobulin domain, a common domain that mediates interactions among immune-regulating molecules (25). U20 is predicted to be a type I membrane glycoprotein possessing a class I MHC-like fold, and as such, we would hypothesize that it traffics through the secretory system to the plasma membrane with cellular receptors involved in immune modulation. To facilitate future functional analysis of U20, we have characterized the expression and trafficking of U20 from HHV-6A and-6B. Here we demonstrate that U20 is a heavily glycosylated type I integral membrane protein that traffics slowly through the secretory system. Additionally, we demonstrate that despite 92% amino acid identity, the steady state localization of HHV-6A and -6B U20 may differ, consistent with their reported different functions (1, 2). Finally, we identify post translational modifications in HHV-6B U20 that might guide future studies with respect to its function.

## Results

### U20 proteins are expressed during roseolovirus infection and are targeted to the secretory pathway

To examine the expression of HHV6B U20 during HHV6B infection, we performed immunoblot analysis of U20 from lysates of HHV6B-infected SupT1 cells using a polyclonal antibody directed against the cytoplasmic tail of U20. We observed a polypeptide migrating at approximately 100 kDa that was absent from lysates of control non-infected SupT1 cells (Figure 2, panel A, lanes 1 and 2).

**Figure 2.**
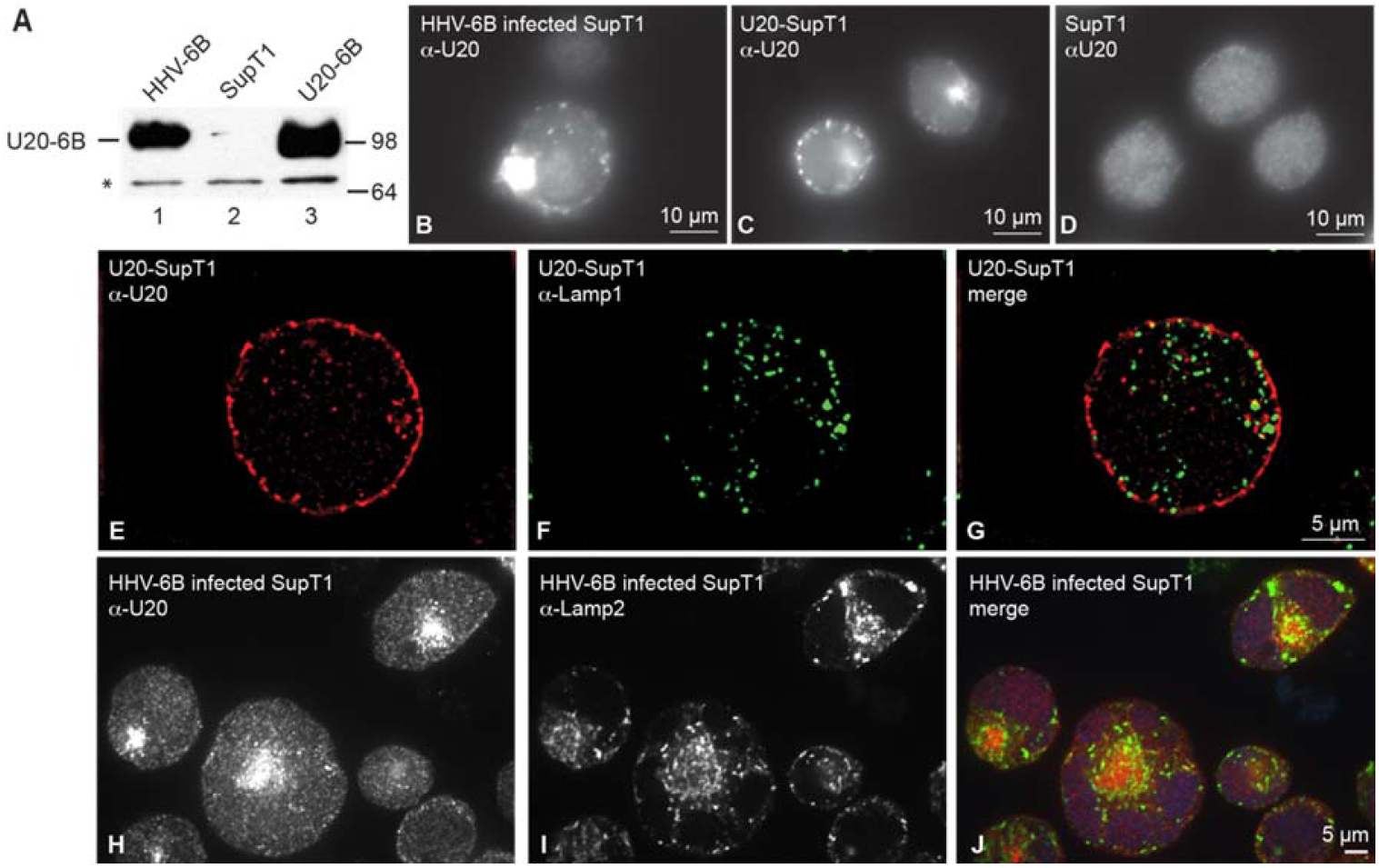
Expression and localization of HHV6B U20 in SupT1 cells. (A) Immunoblot analysis of lysates prepared from HHV-6B infected SupT1 cells or U20-6B expressing SupT1 cells (PPM-U20-6B) using anti-U20 (MCW53). Asterisk indicates a cross-reactive protein recognized by our polyclonal antibody, serving as an internal loading control. (B-D) Immunofluorescence labeling of U20 in B) HHV6B-infected SupT1 cells, C) SupT1 cells stably expressing U20 from HHV6B, or D) or uninfected SupT1 cells. E-G) Structured illumination microscopic co-localization of U20 and lamp1 in SupT1 cells expressing U20. Merged images in (G) illustrates no colocalization. (Pearson’s colocalization coefficient = 0.125). H-J) Confocal immunofluorescence microscopic colocalization of U20 and lamp2 in HHV6B-infected SupT1 cells. Merged image in (J) illustrates no colocalization. Scale bars are shown in bottom right corners of each photoset.

Infection and propagation of HHV6B in culture presents challenges: since the virus is best propagated through cell-cell contact, generation of high titer cell-free HHV6B virus is not possible -nor has a recombinant HHV6B BAC been developed. Thus, to facilitate our study of U20 in a uniformly-expressing cell population, we generated SupT1 cell lines that exogenously express HHV6B U20. We first compared the expression of U20 in these cells with U20 in HHV6B-infected cells. Expression of HHV6B U20 in our stable SupT1 cell line was low, likely due to silencing of the CMV promoter. To enhance the expression level of U20, we incubated the U20 SupT1 cells in phorbol myristate acetate (PMA), which has been shown to enhance expression from the CMV promoter (Fig 3)((26–28)). Exogenously-expressed HHV6B U20 mirrored U20 from HHV6B-infected cells, and also migrated at approximately 100 kDa. Importantly, the level of U20 expression in PMA-stimulated cells was similar to that of HHV6B infected cells. (Figure 2, panel A, lanes 1 and 3).

**Figure 3.**
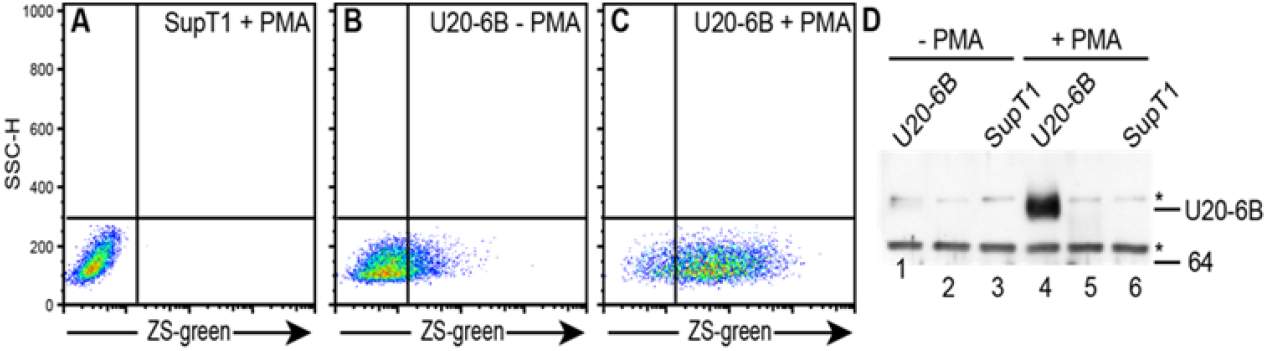
Effect of PMA on CMV promoter driven expression of U20 in SupT1 cells. (A-C) Stimulation of Zs-Green expression in SupT1 cells stably transduced with vector alone (A) or the vector containing the HHV6B U20 coding sequence followed by IRES-driven *Zs-Green* (B and C). Cells in A and C were treated with 40 ng/ml PMA for 20 hours prior to detection of Zs-green by flow cytometry. (D) Immunoblot analysis of RIPA lysates (20µg) prepared from U20-expressing SupT1 cells, probed with a-U20 (MCW53). PMA treatment conditions were as described for A and C.

Immunofluorescence localization of U20 in HHV6B-infected SupT1 cells or U20-expressing SupT1 cells both showed a punctate labeling pattern (Figure 2, panels B and C). The punctate appearance of HHV6B U20 resembled the lysosomal localization of HHV-6 and -7 U21 in T cells (Wang, Q., and Hudson, A., unpublished data). To examine the possibility that HHV6B U20 was localized to lysosomes in HHV6B U20-expressing SupT1 cells, we performed double-label immunofluorescence microscopy using antibodies directed against U20 and lamp1, a marker for the late endosomal/lysosomal pathway (29). HHV6B U20 did not colocalize with lamp1, and instead appeared to localize at the cell surface and in internal puncta (Figure 2, panels E-G). Similar results were seen in HHV6B-infected SupT1 cells using antibodies directed against U20 and lamp2, another marker for late endosomes/lysosomes (Figure 2, panels H-J).

HHV6B U20 shares 91.2% identity with HHV6A U20 (Figure 1). Surprisingly, however, our polyclonal antibodies directed against the cytoplasmic tail HHV6B U20 did not cross-react with HHV6A U20. HHV6A is the only one of the three roseoloviruses for which a recombinant BAC system has been developed, allowing genetic manipulation of the HHV6A genome (30). To examine expression and localization of U20 in HHV6A-infected cells, we generated a recombinant HHV6A virus expressing an U20-mCherry fusion protein. Consistent with the immunofluorescence localization of U20 in HHV6B-infected cells, U20-mCherry in HHV6A-infected cells was also localized in puncta (Figure 4A). In these cells, however, co-localization experiments showed striking co-localization of U20-mCherry with the lysosomal membrane protein lamp2 (Pearson’s colocalization coefficient 0.699) but not with the Golgi membrane protein giantin (Pearson’s colocalization coefficient 0.152)(Figure 4). Thus, HHV6B U20 is localized in puncta distinct from lysosomes or Golgi, whereas HHV6A U20-mCherry colocalizes with lamp2 in a lysosomal compartment.

**Figure 4.**
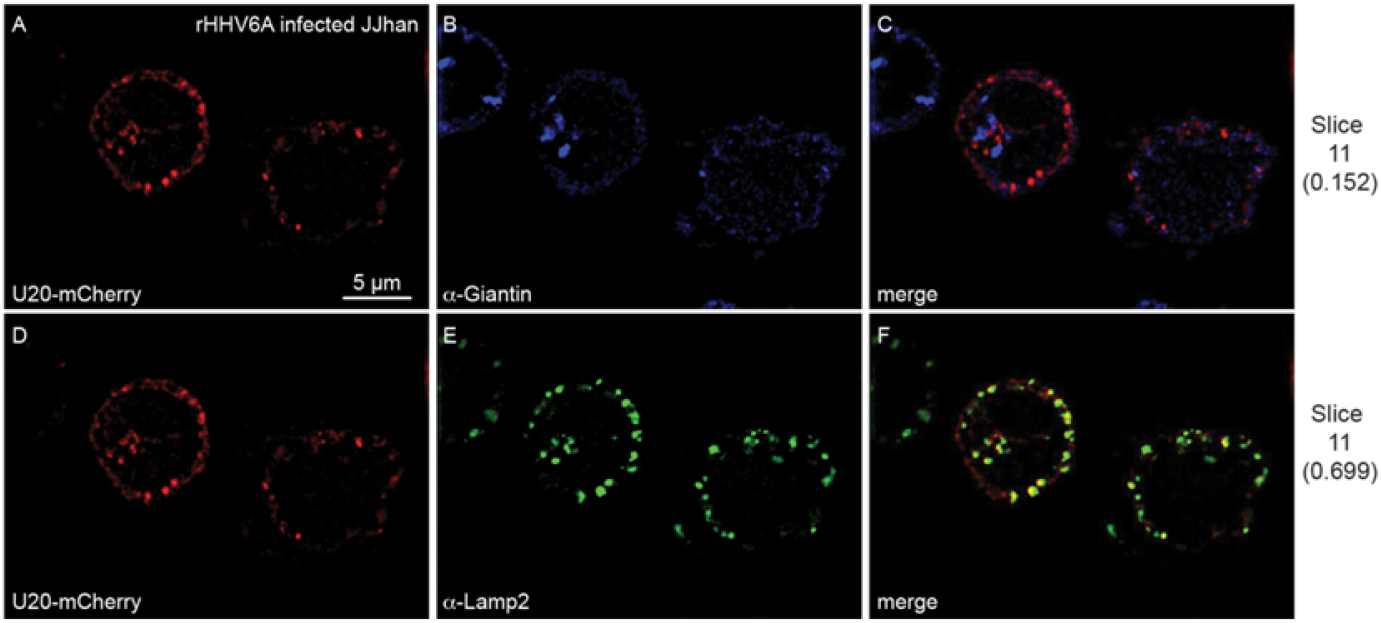
Recombinant HHV6A-infected T cells expressing U20-mCherry fusion. JJhan T cells infected with recombinant HHV-6A, engineered to contain mCherry fused in-frame at the C-terminus of U20. A-C) Confocal still images of rHHV6A U20mCh-infected JJhan cells labeled with anti-Giantin (Golgi marker) and D-F) with anti-lamp2 (lysosomal marker). Pearson’s correlation coefficients are noted to the right. Scale bar, 5 um.

### U20 acquires several post translational modifications as it traffics through the secretory system to the cell surface

Since the localization and SDS-PAGE migration of exogenously-expressed HHV6B U20 mirrored that of U20 in HHV6B-infected cells, we chose to further characterize the trafficking of exogenously-expressed U20. As described above, PMA stimulation was required to achieve physiological levels of HHV6B U20 in our SupT1 cells. Because PMA also activates protein kinase C, which can lead to pleotropic changes in transcription and protein trafficking (31–34), for the biochemical analysis of HHV6B U20, we opted to express U20 in 293T cells. We performed pulse-chase analysis to examine the trafficking of U20 in cells expressing untagged U20, as well as in cells expressing N-terminally tagged U20 and C-terminally tagged U20 (TAP U20-6B and U20-6B TAP, respectively). The TAP tags consists of an HA tag fused to a streptavidin-binding protein (SBP). After a 45-minute pulse label, U20 migrated as an 85 kDa polypeptide (Figure 5, panel A, lane 1), which was converted to an approximately 100 kDa polypeptide at later chase points (Figure 5, panel A, lanes 2-4). N- and C-terminally tagged U20 showed identical results, and reflected the slowed mobility contributed by the TAP tag (Figure 5, panel A, lanes 5-12). The rate of conversion of U20 from the 85kDa form to the 100kDa form is similar for all three U20 molecules (Figure 5, panel B), demonstrating that addition of N-or C-terminal tags to U20 do not affect the kinetics of U20 trafficking.

**Figure 5.**
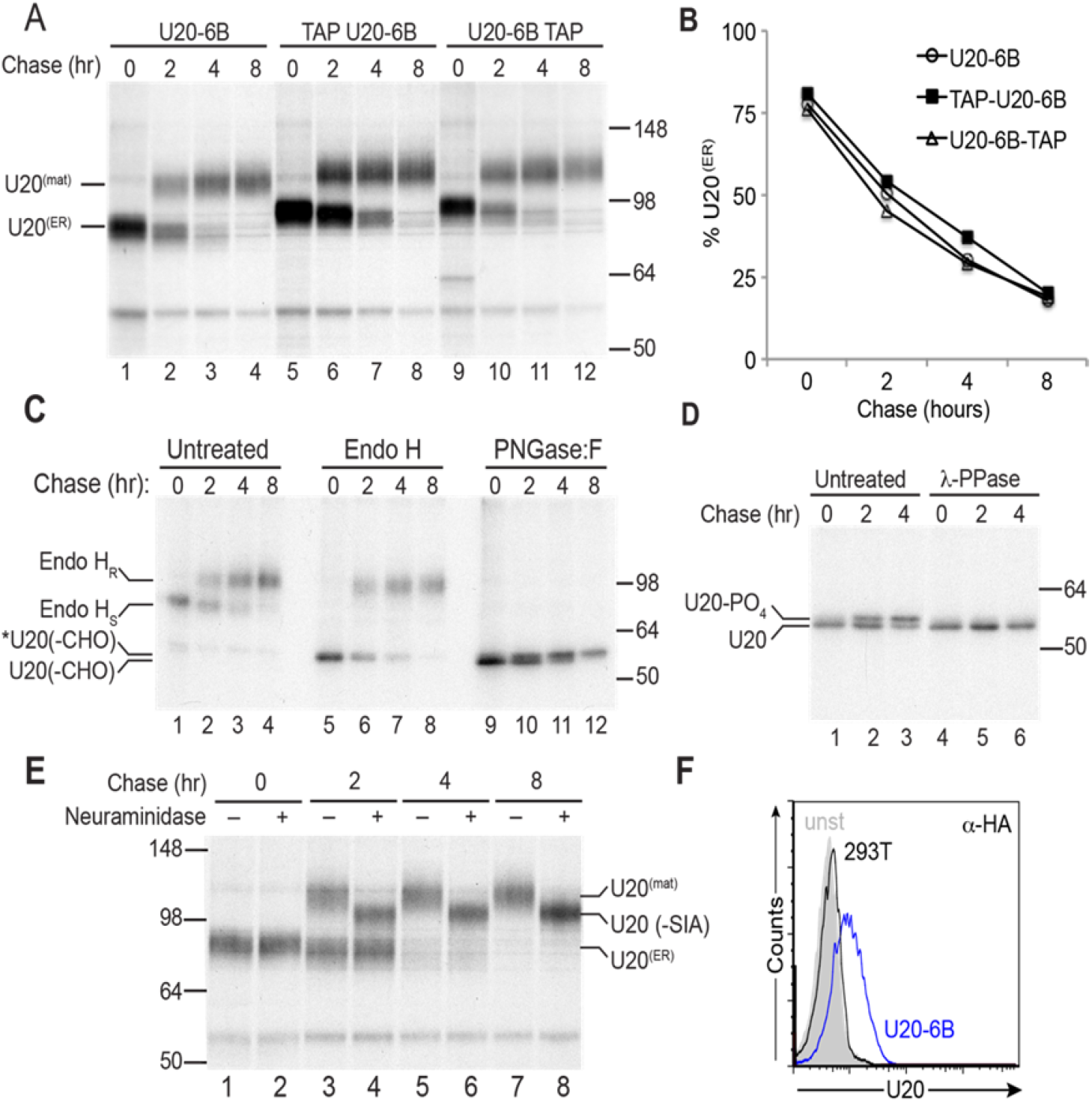
Trafficking and post-translational modifications of HHV6B U20. (A) Pulse chase analysis of TAP-tagged and untagged HHV6B U20 in 293T cells. Cells were pulse-labeled for 45 minutes and chased for 0, 2, 4 or 8 hours. U20 was immunoprecipitated using anti-U20 (MCW53) and gels were analyzed by SDS-PAGE and autoradiography. Migration positions of mature (U20^(mat)^ and immature U20 (U20^(ER)^) are indicated. (B) Quantification of (A) where % U20(^ER^)= 100 x (U20(^ER^)-Bkgrd)/[(U20(^ER^)-Bkgrd) + (U20(^mat^)-Bkgrd)]. (C) Immunoprecipitates from (A) were treated with EndoH or PNGaseF. Migration positions of Endo H resistant (Endo H_R_) or sensitive (EndoH _S_) and fully deglycosylated U20 (U20(-CHO)) are indicated. The slower migrating deglycosylated U20 is indicated by an asterisk. (D) Immunoprecipitations were performed as in (A) and treated with λ-phosphatase followed by PNGase:F and analyzed as above. Phosphorylated (U20-PO4) and dephosphorylated forms of U20 (U20) are indicated. (E) Immunoprecipitations were performed as in (A) and treated with neuraminidase. Migration positions of mature (U20^(mat)^, desialylated (U20 (-SIA)), and immature U20 (U20^(ER)^) are indicated. (F) Flow cytometric analysis of 293T cells expressing an N-terminally tagged HA-U20. Surface expression of U20 was detected with an anti-HA antibody (HA.11). E-G) Structured illumination microscopic immunolocalization of U20 and lamp1 in SupT1 cells expressing U20. Merged image in (G) illustrate no colocalization. H-J) Confocal immunofluorescence microscopic colocalization of U20 and lamp2 in HHV6B-infected SupT1 cells. Merged image in (J) illustrates no colocalization. Scale bars are shown in bottom right corners.

We suspected that the 85 kDa polypeptide corresponds to newly synthesized, ER-resident U20, and that additional modifications to its N-linked glycans acquired upon trafficking through the Golgi are responsible for its shift to 100 kDa. We therefore examined the temporal acquisition of U20 resistance to either Endoglycosidase H (EndoH) or Peptide N-glycosidase:F (PNGase:F). As expected, digestion with EndoH resulted in conversion of the 85 kDa form to approximately 55 kDa, consistent with the 50 kDa predicted molecular weight of U20, thus demonstrating that the 85 kDa form is the ER-localized, newly-synthesized high-mannose form of U20 (Figure 5, panel C, lanes 1 and 5). The 100 kDa form of U20 was resistant to EndoH digestion, confirming that this form of U20 obtained complex-type oligosaccharide modifications in the late Golgi (Figure 5, panel C, lanes 4 and 8). The approximate 30 kDa difference between the EndoH-sensitive and EndoH-resistant forms suggests that all nine predicted N-linked glycan sites are glycosylated. These results suggest that U20 traffics slowly from the ER to Golgi, requiring ∼2 hours for half of the protein to leave the ER.

Interestingly, digestion with PNGAse:F resulted in a doublet with the intensity of the upper band mirroring the intensity of the 100kDa form and the intensity of the lower band mirroring the intensity of the 85kDa form (Figure 5, panel C, lanes 1-4 and 8-12). Thus, although most of the 15 kDa reduction in mobility through the SDS gel is due to addition of complex-type N-linked oligosaccharide modification, an additional post-translational modification that slows the migration of U20 by approximately 1 kDa may occur as the protein traffics beyond the medial Golgi.

Since phosphorylation is a common modification that can result in slightly decreased mobility, we evaluated whether the 100kDa form of U20 was phosphorylated. We treated immunoprecipitates of U20 with λ-protein phosphatase, a manganese-dependent phosphatase with specificity toward phosphorylated serine, threonine, and tyrosine residues. Dephosphorylated proteins were then treated with PNGAse:F to facilitate detection of a reduction in mobility after phosphatase treatment. Interestingly, λ-phosphatase treatment resulted in the disappearance of the upper band of the doublet and a concomitant increase in the intensity of the lower band of the doublet, thus confirming that the glycosylation-independent additional posttranslational modification to U20 is phosphorylation (Figure 5, panel D, lanes 4-6 and 1-3)

Our pulse-chase experiment demonstrates that U20 traffics beyond the medial Golgi. To examine the further trafficking of HHV6B U20, we monitored the rate at which U20 acquired sialic acid residues in the trans Golgi network, assessing sensitivity to neuraminidase digestion. Treatment with neuraminidase did not alter the migration of the 85kDa ER form (Figure 5, panel E, lanes 1 and 2). Neuraminidase treatment did result in increased mobility of the 100kDa form, suggesting that at least some of the nine N-linked glycans become sialylated after they acquire resistance to PNGAse:F (Figure 5, panel E, lane 3-8).

Because U20 has homology to class I MHC heavy chain molecules (25), we hypothesize that U20 may function in some type of immune recognition. As such, we might expect U20 to be present on the cell surface of HHV6B infected cells. Although localization of U20 in SupT1 T cells shows that the majority of HHV-6B U20 localized to punctate compartments distinct from lysosomes at steady-state, we surmised that some of the protein might reach the cell surface. Since our TAP-tagged U20 proteins traffic similarly to untagged U20 molecules, we employed flow cytometry of non-permeabilized TAP-U20-expressing cells to determine whether N-terminally tagged U20 appeared at the cell surface. Flow cytometry results demonstrated that some U20 is present on the cell surface (Figure 5, panel F).

To examine the presence of U20 on surface of HHV6B-infected cells, we generated an antibody directed against the extracellular domain of U20. Flow cytometry results using this antibody were consistent with our results in 293T cells expressing HA-tagged U20, showing that some U20 is detectable on the surface of HHV6B-infected SupT1 cells (Figure 6). The amount of U20 on the surface of both cell types seemed low relative to the overall expression level of U20, and steady-state immunofluorescence experiments did not reflect a strong cell surface presence of U20 (Figure 2, panels B and H). We therefore hypothesized that perhaps U20 reaches the cell surface and is rapidly internalized, reducing the amount of U20 seen on the surface at steady-state.

**Figure 6.**
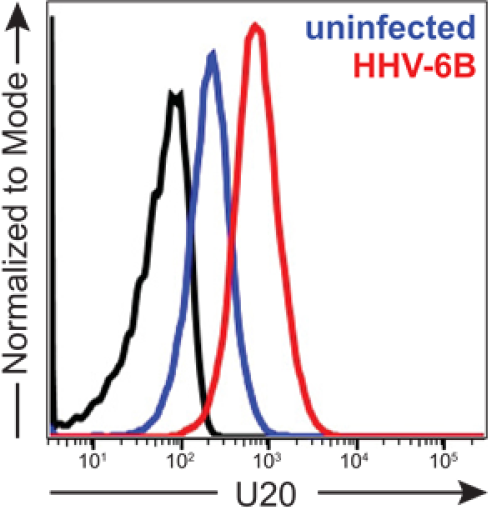
Surface presence of HHV6B U20. (A) Flow cytometry of non-permeabilized HHV6B-infected (red) and noninfected (blue) SupT1 T cells incubated with anti-U20 Black trace: no primary Ab.

To examine whether U20 is internalized after reaching the cell surface, we incubated HHV6B-infected cells with extracellularly-or intracellularly-directed U20 antibodies at 4°C, and then warmed the cells to 37°C to allow the cells to internalize the antibody complexed to U20. We simultaneously labeled the cells with an antibody directed against the extracellular portion of class I MHC molecules, which normally localize to the cell surface. Surface-expressed U20 decorated with the extracellularly-directed U20 antibody MCW64 was localized to punctate compartments following a 30-minute internalization at 37°C (Figure 7, panel A), while HHV6B-infected cells incubated with the antibody directed against the cytoplasmic tail of U20 showed no internalization of the Ab (MCW53, Figure 7 panel C). Interestingly, cells showing the most internalized U20 exhibited very little class I MHC labeling (Figure 7A-B, pink arrows), while cells showing very little internalized U20 exhibited prominent surface class I MHC (Figure 7A-B, white arrowheads).

**Figure 7.**
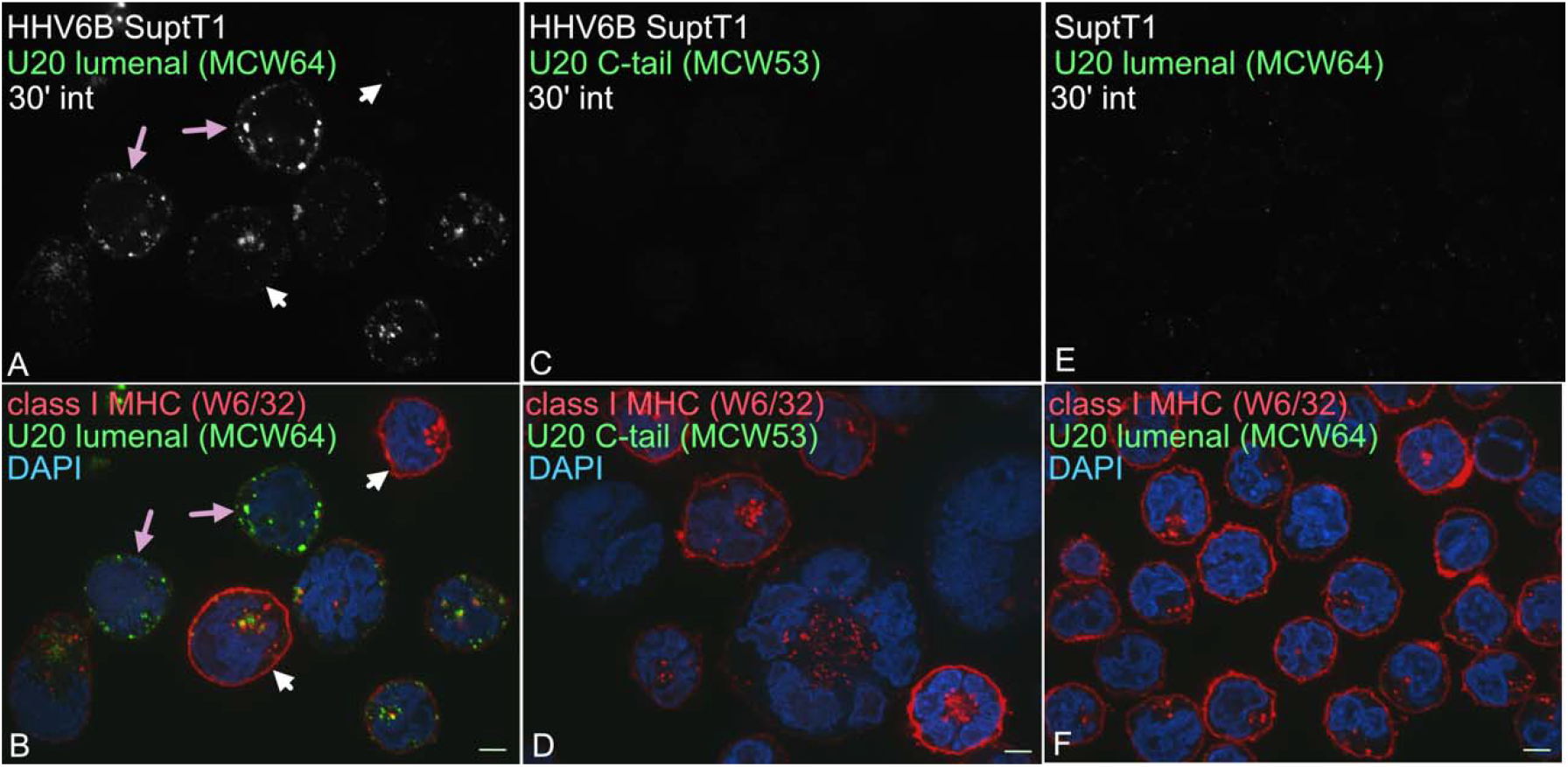
Trafficking of HHV6B U20 to the cell surface. (A-F) HHV6B-infected (A-D) or control SupT1 cells (E-F) were incubated with antibodies directed against the either the extracellular portion of class I MHC molecules (W6/32) and antibodies directed against the extracellular (MCW64)(A,B) or intracellular (MCW53) portions of U20 (C,D) on ice. Cells were then warmed to 37°C for 30 minutes before fixation, permeabilization, and incubation with AlexaFluor-conjugated secondary antibodies. White arrowheads denote cells with high expression of class I MHC molecules. Pink arrows denote cells in panel A with internalized MCW64 Ab. Surface class I MHC molecules are absent in these cells (panel B). Scale bars, 5um.

Class I MHC molecules are rerouted to lysosomes for degradation by the HHV6B U21 gene product. We think it likely that the cells that exhibit the most internalized U20 are HHV6B-infected cells that also express more U21 (15)(35). In this asynchronously infected cell population, we estimate that HHV6B infects approximately 50-70% of the cells; cells expressing high levels of surface class I MHC molecules may either not yet be infected with HHV6B or may be at an early stage of infection not yet resulting in degradation of class I MHC molecules. As expected, noninfected SupT1 cells labeled with MCW64 showed little to no internalization and uniformly high labeling of class I MHC molecules (Figure 7,E-F). Collectively, these results demonstrate that U20 is a type I integral membrane protein that traffics slowly through the secretory pathway to the cell surface, where it is then rapidly internalized.

## Discussion

The *U20* gene lies within a cluster of Roseolovirus-specific genes (*U20-U24*) that are dispensable for viral growth *in vitro*, suggesting a possible role in host evasion (36). Within this cluster, the U20 and U21 open reading frames encode proteins with predicted homology to class I MHC-like protein folds (37)(25). HHV-6A, -6B, and -7 U21 proteins all function to redirect class I MHC molecules to lysosomes, and HHV-6A, -6B, and -7 U24 proteins all function to downregulate the T cell receptor CD3 from the cell surface (20). Interestingly, HHV6A U20 and HHV6B U20 share 92% amino acid identity (Figure 1) yet results from two published studies suggest different functions for U20 from HHV-6A and -6B (1, 2): HHV6A U20 expression was shown to downregulate NKG2D ligands, while HHV6B U20 expression was shown to inhibit TNFα-induced apoptosis during nonproductive infection with HHV6B (1, 2).

Several studies have examined the kinetics of HHV6A gene expression during infection (23, 24). Although these studies indicate that the *U20* gene is transcribed during infection, to our knowledge, ours is the first study to demonstrate the U20 protein is expressed during infection. Using a polyclonal antibody directed against the cytoplasmic tail of HHV6B U20, we detected an approximately 100 kDa polypeptide in lysates from HHV6B-infected SupT1 cells (Figure 2). Given the similar size of this polypeptide to an exogenously expressed U20, we conclude that the full-length protein as annotated is expressed in infected T cells. We demonstrate here that all nine possible N-linked glycan sites present within HHV6B U20 ORF appear to be used, consistent with recent structural predictions (25). As U20 traffics through the Golgi compartment, these high-mannose N-linked glycans receive complex-type modifications, including sialylation, resulting in an additional ∼15 kDa in size (Figure 5). Interestingly, this mature form of U20 is also phosphorylated on one or more serine, threonine, or tyrosine residues. The cytoplasmic tail of HHV-6B U20 has eight predicted phosphorylation sites while HHV-6A has five, and four of them are conserved in both proteins (Figure 8).The role of phosphorylation, if any, in regulating trafficking or the function of U20 remains to be investigated.

**Figure 8.**
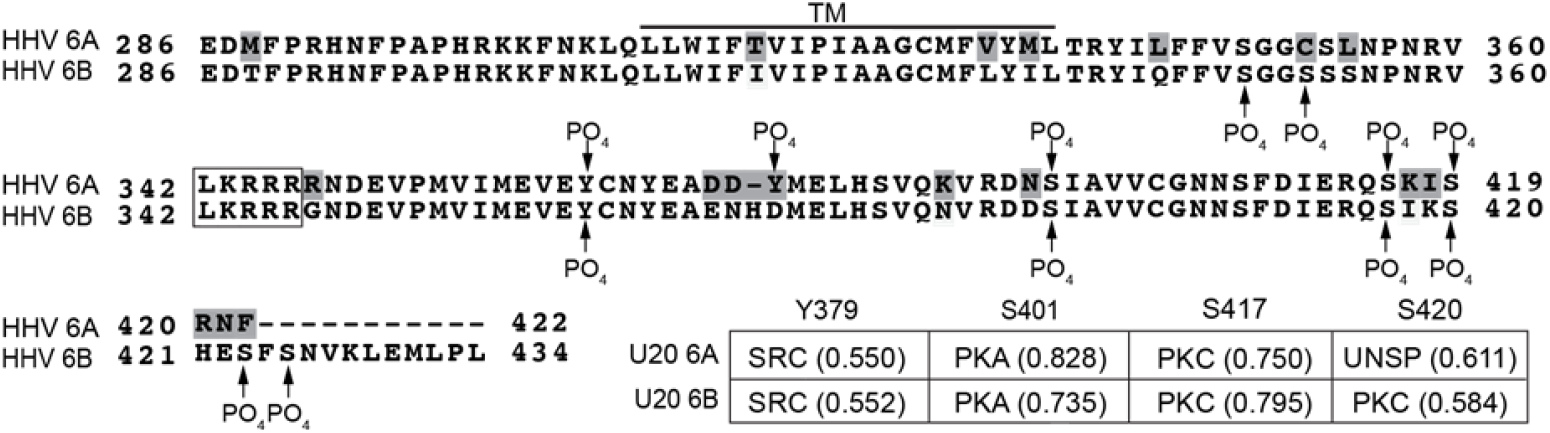
Alignment of U20 from HHV6A and-6B including possible phosphorylation sites. Alignments in Figure 1 was modified to include phosphorylation sites in U20 6B as predicted by NetPhos 3.1. Only sites with scores higher than 0.5 are indicated. Sites predicted to be phosphorylated in HHV6A U20 are indicated above the sequence and those for HHV6B U20 are below. Inset table shows the top predicted kinase for each of the four phosphorylation sites conserved in both proteins. Numbering of the amino acid is based on that of HHV6B U20, and the scores are in parentheses after the kinase name. A potential di-arginine retention motif is boxed.

U20 is a type I membrane glycoprotein and, as such, it should localize somewhere within the secretory or endomembrane system. Immunofluorescence analysis of HHV6A- and 6B-infected cells showed that HHV6A U20 was localized to lysosomes and that HHV6B U20 was localized to the cell surface and to puncta within the cell. The HHV-6A U20-mCherry puncta colocalized with the lysosomal marker lamp2 during infection, and the HHV-6B U20 puncta did not. It is possible that fusion of mCherry to the C-terminus of HHV6A U20 in some way altered its localization, but without an antibody to native HHV6A U20, we were unable to investigate this possibility. We note that a similarly placed C-terminal fusion of streptavidin binding protein and an HA moiety to the C-terminus of HHV6B U20 did not alter the trafficking of HHV6B U20 (Figure 5).

Finally, our trafficking studies demonstrate that U20 moves quite slowly through the secretory system. For comparison, approximately 75% of class I MHC heavy chains acquire Endo H resistance after a 30-minute chase period, but newly synthesized U20 is only 50% Endo H-resistant after a 2-hour chase (Figure 5)(18). ER retention motifs for type I membrane proteins, including di-lysine (KKxx or xKxK) and di-arginine (RR or RXR) motifs located in the cytoplasmic tail can help retain proteins in the ER, either by preventing their export or by facilitating their retrieval by the COPI machinery (for review, see (38)). While U20 does not appear to contain a di-lysine motif, there is an LKRRRR sequence in the cytoplasmic tail of HHV6A and 6B U20 that might serve as an ER-retention signal (Figure 6)(38). Whether the LKRRR sequence in the cytoplasmic tail is a *bona fide* di-arginine motif remains to be determined.

We hypothesize that the slow trafficking of U20 could be a combination of several factors: U20’s 20-amino acid transmembrane domain is more similar to mean hydrophobic lengths of ER and Golgi localized proteins than plasma membrane resident proteins (20.3 and 20.6 vs 24.4aa) which may decrease the amount of U20 that localizes to ER exit sites, reducing ER export (39). Additionally, since U20 lacks conventional export motifs, it might also experience futile recycling, further delaying trafficking to the Golgi (40, 41). As no binding partners have yet been identified for U20, we have not determined whether interaction with a cellular protein, possibly through its predicted Ig domain, might alter the kinetics of U20 trafficking.

## Materials and methods

### Cells and Reagents

HEK293T cells (ATCC) were cultured in Dulbecco’s modified Eagle medium (DMEM) (Thermo-Fisher LifeTechnologies, Waltham, MA) supplemented with 5% newborn calf serum (NCS) (Thermo-Fisher) and 5% fetal bovine serum (FBS) (LifeTechnologies) and puromycin (400ng/ml) as indicated. SupT1 (immature human T lymphocytic cell) cells were cultured in RMPI supplemented with 10% FBS and puromycin (200ng/ml) as indicated. All cells were grown at 37°C, 5% CO_2_. When indicated, T lymphocytes were stimulated with 40ng/ml phorbol myristate acetate (PMA) for 20-24 hours to increase expression from the CMV promoter (see Figure 3). Chemicals were purchased from Sigma-Aldrich (St. Louis, MO), unless otherwise noted.

### HHV6A and -6B virus propagation

HHV6B Strain Z29, obtained from Dr. Phil Pellet, (CDC, Atlanta, GA) was propagated by co-culturing infected cells with uninfected SupT1 cells at ratios varying from 1:1 to 1:4 (infected:uninfected) in RPMI supplemented with 5% heat inactivated FBS, 100 units/ml penicillin, and 100 µg/ml streptomycin (Thermo-Fisher). Cells were monitored visually for signs of cytomegaly and were typically subcultured or used for experiments 5 to 12 days after incubating with infected cells when cytopathic effects (CPE) was present in 60-80% of the cells.

Wild-type rHHV6A GFP+ BAC (generously provided by Dr. Yasuko Mori (Kobe, Japan)) and rHHV6A GFP+-U20-mCherry viruses used in this study were derived from HHV6A strain U1102, in which the BAC sequence, containing GFP, was inserted between the U53 and U54 polyA signal sequences in the HHV6A genome (30). HHV6A BAC DNA was electroporated into JJhan T cells using Lonza (NEB, New Brunswick, MA). When the cells in the transfected culture reached 80% GFP+, cells were co-cultured with uninfected cells at an approximate ratio of 2:1 (uninfected:infected). HHV6A-infected and noninfected JJhan cells were cultured in RPMI with 5% FBS.

The U20-mCherry fusion protein in HHV6A was constructed using a two-step RED recombination protocol (42). The plasmid pEPmCherry-in (generously provided by Greg Smith (Northwestern University)) was used as a template for PCR to insert mCherry as a translational fusion to the C-terminal end of the HHV6A U20 ORF (43). Primers used for this construct were 5’-<u>aataacctcacagtcatggaaatattttccaaaggtaaaatttctaacttcacattg</u>**cta**CTTGTACAGCTCGTCCATGC-3’and 5’-gttgtctgcggaaataattcgtttgatatagaacgacagtctaaaatctcacgaaattttATGGTGAGCAAGGGCGAGGA-3’. The primers encoded the mCherry sequence to be amplified (upper case), a stop codon (bold), as well as homology to U20 (lower case), and the genomic sequence of HHV6A downstream of U20 (underlined).

### Antibodies

Polyclonal U20 specific antibodies were generated in rabbits using a bacterially expressed and purified GST fusion protein containing an N-terminal GST fused to the cytoplasmic tail of U20 from HHV6B. GST fusion protein purification is detailed below. Purified, native, GST fusion proteins were sent to Cocalico Biologicals (Reamstown, PA) to generate antibodies in rabbits (MCW53). For MCW64, the extracellular (α1 and α2) domain of U20’s class I MHC fold were tagged with 6XHis, purified on a Nickel-NTA column, and provided to Cocalico Biologicals in native form. Antibodies to the HA epitope (HA.11 MMS-101R) or Giantin (PRB-114C) were purchased from Covance (Princeton, NJ). The lamp1 monoclonal (H4A3) and lamp2 monoclonal (H4B4) hybridoma supernatants were generously provided by Dr. Tom August). The secondary goat F(ab’)2 anti-mouse IgG-PE antibody used for flow cytometry was purchased from R&D Systems (F0102B) (Minneapolis, MN). The secondary antibodies used for immunofluorescence were Alexa Fluor 488 or 594 conjugated antibodies from Life Technologies (Thermo-Fisher).

### U20 constructs

U20 from HHV-6B was cloned into lentiviral vectors that were adapted in our laboratory from pHAGE vectors using standard methods (44). Untagged HHV6B U20 contains a single substitution within its signal sequence (I2V) to create an ideal Kozak consensus sequence. TAP-tagged U20s were first cloned into pcDNA3-based vectors that we had modified to contain either an N-terminal cleavable signal sequence and tag, or a C-terminal tag and stop codon, and then subcloned into lentiviral pHAGE vectors. N-terminally TAP tagged U20 contains the signal sequence from the mouse MHC class I molecule H-2Kb (KbSS) followed by the TAP tag (YPYDVPDYAGLNENLYFQGAGTWSHPQFEK) and U20 lacking its predicted signal sequence. C-terminally-tagged U20 contains U20 with the substitution in its normal signal sequence listed above, and the TAP tag, in the reverse orientation, appended to the C terminus.

### Lentiviral transductions

Cell lines were transduced with a lentiviral vector pHAGE-puro-MCS (PPM) in which TAP-tagged or untagged versions of HHV6B U20 was expressed under the control of the CMV promoter, and an IRES-driven puromycin N-acetyl transferase gene (*Pac*) that allowed for puromycin selection. In some instances (Figure 3), SupT1 cells were transduced with a related lentiviral vector pHAGE-MCS (PMG) in which the *Pac* gene was replaced with the gene for Zs-Green. Packaging, envelope, and vector plasmids were cotransfected into HEK293T cells using TransIT®-293 (Mirus Bio, Madison, WI). Lentiviral supernatants were harvested at 48-72 hrs, filtered and either used to infect desired cell lines directly (293T cells) or concentrated prior to infections (SupT1 cells). Lentiviruses were concentrated by ultracentrifugation for 3 hrs at 4°C at 50,000xg, and SupT1 cells were infected on two consecutive days by spinoculation (1000 X g for 2 hr at 30°C) with concentrated viruses. For puromycin-resistant constructs, cells were cultured in selective medium for at least 10 days (400ng/ml puromycin for 293T and 200ng/ml for SupT1). Non-adherent cells transduced with PMG based vectors were typically sorted for top 5-20% ZS-green expressing cells once on a FACSAria III (BD Biosciences).

### Pulse-chase experiments

Cells were detached with trypsin and incubated in methionine- and cysteine-free DMEM (Life Technologies) supplemented with 2% FBS for 30 min at 37°C (starve). The cells were labeled with 250-1000 µCi/ml of [^35^S]-Express label (1100 Ci/mmol; PerkinElmer, Boston, MA) at 37°C depending on the length of the pulse, and chased with complete DMEM supplemented with 1 mM non-radioactive methionine and cysteine for indicated times at 37°C. Cells were washed with PBS then lysed in the buffer indicated. Lysates were centrifuged for 10 min at 16,000 X g at 4°C to pellet nuclei and debris. Clarified lysates were typically incubated overnight at 4°C with designated antibodies, followed by addition of Protein A agarose (Repligen Corporation, Waltham, MA) or protein G agarose (Thermo-Fisher) and further incubated for 1 hour at 4°C. Immunoprecipitates were washed four times with the appropriate wash buffer, and subjected to SDS-PAGE gel electrophoresis. Digestion with Endo H or PNGase F (NEB, Beverly, MA) were performed according to manufacturer’s suggested protocol. Quantification, as indicated, was performed from phosphorimages generated on a Storm 820 (GE Healthcare, Piscataway, NJ) using ImageQuantTL software. Formulas used for normalization varied and are indicated in the figure legends.

### Immunofluorescence microscopy

T cells were adhered onto poly-L-lysine-coated glass coverslips and washed with PBS, fixed with 4% paraformaldehyde in PBS, permeabilized with 0.5% saponin in PBS and 3% BSA, incubated with primary antibodies, washed and incubated with Alexa Fluor 488-or 594-conjugated secondary antibodies (Life Technologies). For immunofluorescence microscopy analysis of U20-mCherry and lamp1, T cells were adhered onto poly-L-lysine-coated glass coverslips and permeabilized in 0.5% saponin in PBS, 3% BSA, 880mM CaCl_2_, and 490 mM MgCl_2_. Permeabilized cells were incubated with primary antibody, washed, and then incubated with secondary antibody conjugated to a fluorophore. Pearson’s correlations were determined in three separate slices from three different images using NIS Elements software.

### Antibody Internalization

HHV6B-infected SupT1 cells were incubated with either MCW53(cytoplasmic U20) or MCW64(extracellular U20) and W6/32 (class I MHC) at 4°C for 30’ in RPMI + 20 mM Hepes, and then warmed to 37° for 30’ to allow internalization of bound antibodies. Cells were washed with cold RPMI, fixed with 4% paraformaldehyde, and incubated with AlexaFluor 488 or 647-conjugated secondary antibodies in 0.5% saponin + 3% BSA, washed, and confocally imaged in suspension at 100X in a 96 well glass-bottomed plate.

### Microscopy

Superresolution microscopy was performed on a Nikon Structured-Illumination Microscopy (N-SIM; Nikon) and NIS Elements AR imaging and 3D reconstruction software (v. 5.11). SIM images were taken using a heated Nikon 100X oil-immersion lens (CFI Apo SR TIRF 1.49 NA) and an Andor iXon+897 EMCCD camera. Confocal microscopy was performed on a Nikon EclipseTi2 microscope equipped with a W1 spinning disc, Orca Flash CMOS camera, and 60X oil-immersion objective (CFI Plan Apo λ 1.4 NA) and NIS-Elements AR imaging and 3D reconstruction software (v. 6.0). Epifluorescence microscopy was performed on a Nikon Eclipse TE2000-U microscope with Metamorph Software (v. 7.0r4) equipped with a Photometrics Coolsnap EZ camera and 60X oil-immersion objective (CFI Plan Apo VC 1.4 NA).

### Flow cytometry

Adherent 293T cells were detached with trypsin in PBS prior to labeling. Cells were washed in ice-cold PBS and incubated with the HA.11 monoclonal antibody in 1% BSA/PBS for 45 min on ice. The cells were then washed with 1% BSA/PBS and incubated with goat F(ab’)2 anti-mouse IgG-PE (F0102B) in 1% BSA/PBS for 45 min on ice. Flow cytometry was performed on a Guava Easycyte mini (Millipore, Billerica, MA) and the data was analyzed using FlowJo analysis software (Treestar, Ashland, OR). For HHV6B-infected cells, cells were incubated with primary antibodies in 1% bovine serum albumin (BSA) in phenol red-free DMEM for 30 min on ice, washed, and incubated with secondary antibodies. Flow cytometry was performed using an LSRII flow cytometer (BD Biosciences, San Jose, CA), and data was analyzed using FlowJo analysis software (v. 10.7, BD Biosciences). Non-viable cells were excluded from all flow cytometric analyses.

### Immunoblot analysis

Cell lysates were prepared in RIPA buffer (50 mM Tris-HCl [pH 7.4], 150 mM NaCl, 1% Triton X-100, 1% sodium deoxycholate, 0.1% SDS, 1 mM EDTA). Lysates were normalized to total protein concentration as determined by BCA assay (Thermo-Fisher). Lysates were resolved by SDS-PAGE electrophoresis, transferred to BA-85 nitrocellulose membrane (Millipore-Sigma, Florham Park, NJ) and probed with designated primary antibodies followed by a goat anti-rabbit IgG HRP-conjugated secondary antibody (BioRad, Hercules, CA). Bands were visualized using SuperSignal reagent (Thermo-Fisher) and quantified with an Alpha Imager (ProteinSimple, San Jose, CA).

## Acknowledgements

We thank Drs. Grant Weaver and Larry Stern for helpful discussion, and Colin Parsons for excellent technical assistance. We acknowledge funding from National institutes of Health under award numbers AI076722 and S1 GM120735 to A.W.H.

